# Antimicrobial activity of flavonoids glycosides and pyrrolizidine alkaloids from propolis of Scaptotrigona aff. postica

**DOI:** 10.1101/2021.07.01.450350

**Authors:** TM Cantero, PI da Silva Junior, Giuseppina Negri, RM Nascimento, RZ Mendonça

## Abstract

Stingless bees belonging to the Meliponinae subfamily, are known as meliponines. Scaptotrigona affinis postica Latreille, 1807 from northeast of Brazil is popularly known as ‘tubi’ in Maranhão State. Scaptotrigona, which is widely distributed in neotropical regions, includes species that build their hives in pre-existing cavities. Flavones di-C-glycosides, and the pyrrolizidine alkaloid 7-(3-methoxy-2-methylbutyryl)-9-echimidinylretronecine were reported previously in propolis from S. postica. Fractions 40 AEP and 40 MEP from ethanolic extract were analyzed by LC-MS. The chromatographic profile of fractions 40 AEP and 40 MEP revealed the presence of many pyrrolizidine alkaloids, among them, lithosenine (**14**), lithosenine arabinoside (**19**), 7-angeloyl-9-(2,3- dihydroxybutyryl) retronecine (**1**), 7-(2-methylbutyryl) retronecine (**3**), 9-sarracinoylretronecine (**13**) and viridinatine (**8**),besides the flavonoids schaftoside (**15**), aromadendrin-7-O-methyl ether (**12**), 7-methoxy-5,6,3’,4’,5’,6’-hexahydroxy-flavone-3-O-glucuronide (**11**), mangiferin (**10**) and 3-O-methyl mangiferin (**17**). Fractions 40 AEP and 40 MEP showed antimicrobial activity against Gram negative bacteria, including Escherichia coli D31-streptomycin resistant. Cell viability was expressed in terms of the relative absorbance of treated and untreated cells (control). There was no statistical difference between treated and untreated cells.

## Introduction

Bees (Hymenoptera: Apoidea) and stingless bees (Apiade, Meliponini, Meliponinae subfamily) are important pollinating agents with great importance for plant diversity. The Meliponinae subfamily is divided into two tribes: Meliponini and Trigonini. In the North and Northeast regions of Brazil, there is a great diversity of native stingless bees with large number of species (Pedro, 2014). In the state of Maranhão, northeast of Brazil, the most relevant species are Melipona fasciculata Smith 1858, popularly known as tiúba, and Scaptotrigona affinis postica Latreille, 1807, popularly known as tubi. Scaptotrigona genus is widely distributed in neotropical regions and includes species that build their hives in pre-existing cavities (Lopes et al., 2020; Pedro, 2014, Araújo et al., 2011). As honeybees, stingless bee produce honey, pollen, propolis or geopropolis. they possess stingers, but its cannot be used for their defense (Lavinas et al., 2019; Lopes et al., 2020; Ferreira et al., 2020). Stingless bee products are promising sources of biologically active compounds, because are collected in the tropical and subtropical regions of world, where exist a rich vegetation (Al Hatamleh et al., 2020). Stingless bee honey exhibited antimicrobial activity against Gram-positive bacteria: Bacillus subtilis, Micrococcus luteus, Bacillus megaterium and Bacillus brevis, as well as Gram-negative bacteria: Escherichia coli and Pseudomonas syringae (Al-Hatamleh et al., 2020).

Some stingless bee species produce geopropolis, which contain soil into resinous material (Pereira et al., 2020). Propolis and geopropolis exert many functions, among them, to protect beehives against insects and pathogenic microorganisms, to restrict entry into the hive, to line the interior walls of the hive, to strengthen the honeycombs, and embalm animals (Lavinas et al., 2019; Tran et al., 2020). Propolis is produced with the resin that bees collect from the cracks in the bark and leaf buds of different trees, including different species of poplars, conifers (pines and cypress), birches, alders, willows, palms, chestnuts, eucalyptus, acacia, Clusia spp., and Baccharis dracunculifolia DC. (Šturm and Ulrih, 2020; Kocot et al., 2018; Anjum et al., 2019; Pasupuleti et al., 2017). Their chemical composition varies according to the bee species, flora visited by bees, seasonality and the region of collection (Letullier et al, 2019; Ferreira et al.,2020; de Mello Sousa et al. 2019). In some studies were observed similar chemical profile for different propolis samples from *Apis mellifera* L. and propolis or geopropolis samples from stingless bee, indicating that bees actively and selectively forage resins source containing bioactive compounds, with antimicrobial (antibacterial, antifungal, antiparasitic, and antiviral properties), in order to protect themselves against pathogens and predators, suggesting that both groups of bees can collect the same resin source (Ferreira et al., 2017a,b, Tran et al., 2020; Lavinas et al., 2019). Quercetin methyl ethers, and methoxy chalcones, similar than that detected in young and adult leaflets of Mimosa tenuiflora, were detected in propolis from Apis mellifera and geopropolis from Scaptotrigona aff. depilis, indicating Mimosa tenuiflora as probably resin source (Ferreira et al., 2017a,b). Flavonoid aglycones (mainly neoflavonoids, isoflavonoids) and pterocarpans were detected in extracts of propolis from Apis mellifera and Trigona sp (Okińczyc et al., 2020).

More than 600 compounds were isolated from propolis until 2018 (Šturm and Ulrih, 2020; Anjum et al., 2019; Tran et al., 2020, Lavinas et al., 2019). The high number of different chemical classes of compounds provide to the propolis a bioactive potential, with scavenging characteristics that is important to avoid free radical damages in the human health (Ferreira et al., 2020, Tran et al., 2020, Lavinas et al., 2019). In stingless bee propolis or geopropolis were found phenolic acids, hydrolysable tannins (de Sousa-Fontoura et al., 2020; Dutra et al. 2014, Torres et al., 2018), pentacyclic triterpenes and cycloartane-type triterpenes (Yam-Puc et al., 2018, Pujirahayu et al., 2019a,b), diterpenic acids (Carneiro et al., 2016, Sawaya et al., 2009), alkylphenols (Negri et al., 2019), fatty acids, ga**ll**oyl glucosides, e**ll**agic acid, acyl–hexosides, acyl- ga**ll**oyl- hexosides (Souza et al., 2018, de Sousa Fontoura et al., 2020), flavonoids (Torres et al., 2018, de Sousa Fontoura et al., 2020), flavonols methyl ethers and methoxy chalcones (Ferreira et al., 2017a,b), phenylpropanoids (de Sousa-Fontoura et al., 2020), steroids and saponins (Ibrahim et al., 2016) and catechins and epicatechin (Hochhein et al., 2019). Flavonoids glycosides were detected in propolis for the first time between 2013 and 2018 (Righi et al., 2013). Flavones di-C-glycosides, flavones C-glycosides and alkaloids were found in propolis from Scaptotrigona affinis postica bees (Coelho et al., 2015, 2018), in geopropolis from Scaptotrigona bipunctata bees (Cisilotto et al., 2018) and from Algerian propolis (Soltani et al., 2017). Flavonoids, derivatives of glycosylated phenolic acids and terpenoids were detected in hydroalcoholic extract from Melipona orbignyi (Dos Santos et al., 2017a). Catechin, epicatechin, gallocatechin, epicatechin gallate, aromadendrin, naringenin, pinocembrin, p-coumaric acid (Hochhein et al., 2019), quercetin, p-hydroxy-benzoic acid, caffeic acid, coumaric acid, gallocatechin, rutin, gallic acid and syringic acid were detected in propolis extract from Melipona quadrifasciata and propolis from Tetragonisca angustula (Dos Santos et al., 2017b)

Propolis samples from Tetragonisca angustula (Carneiro et al., 2016) and from Scaptotrigona species (Sawaya et al., 2009) independently of their geographic origin, presented diterpenic and triterpenic acids similar than that detected in flowers from Schinus terebinthifolius Raddi (Anacardiaceae), their probable resin source. On the other hand, quercetin methyl ethers, and methoxy chalcones were detected in propolis from Scaptotrigona aff. depilis collected in the state of Rio Grande do Norte, Brazil (Ferreira et al., 2017b), while stigmasterol, taraxasterol, vanilic acid, caffeic acid, quercetin, luteolin, and apigenin were detected in propolis sample from Scaptotrigona depilis collected in the state of Mato Grosso do Sul, Brazil (Bonamigo et al., 2017). Piperidine alkaloids together flavones-C-glycosides were detected in geopropolis from S. bipunctata, while geopropolis from Melipona quadrifasciata collected in the same locality, Florianópolis, Santa Catarina, Brazil, were detected terpenoids and flavonoids (Cisilotto et al., 2018).

Melissopalynological studies focusing on pollen grains found in the sediments of products elaborated by bees, can help determine their botanical and geographical origins. In the northeast region of Brazil, the pollen types most collected by bees were from Borreria verticillata, Croton, Cecropia, Eucalyptus, Mikania, Mimosa caesalpiniifolia, Myrcia, Poaceae type, Solanum and Schinus, while in the southeast, the pollen types most collected were from Anadenanthera, Cecropia, Citrus, Eucalyptus, Eupatorium, Mimosa caesalpiniifolia, Myrcia and Poaceae type (de Souza et al., 2019). Gluconic acid, quercetin-3,4-diglucoside, apigenin-6-*C*-glucoside, isoorientin-2”-*O*-rhamnoside, kaempferol 3,7-di-*O*-rhamnoside and ellagic acid were detected in pollen extract collected from Scaptotrigona aff. postica, in Maranhão state, Brazil (Lopes et al., 2020).

Propolis and geopropolis exhibit antiseptic, antioxidant, antibacterial, antifungal, antiviral, antiparasitic, hepatoprotective, immunomodulatory and anti-cancer activities, besides to be used in the treatment of dermatoses and other diseases (Letullier et al, 2019, Santos et al., 2020, Ferreira et al.,2020, Lavinas et al., 2019, Tran et al., 2020, Anjum et al., 2019, Ali and Kunugi, 2020). Ethanolic extracts of propolis from Scaptotrigona depilis and Melipona quadrifasciata anthidioides showed cytotoxic activity against erythroleukemic cells (Bonamigo et al., 2017; Dos Santos et al., 2017d).

Propolis samples from Scaptotrigona sp collected in the Serra do Corda region (Maranhão State, Brazil) caused a decrease in colony formation in glioblastoma cell lines and had no effects on apoptosis, demonstrating a cytostatic action (Borges et al., 2011). Propolis from S. postica is used by the population of Barra do Corda, Maranhão state, for the treatment of tumors and wound healing (Araújo et al., 2010; Lopes et al., 2020), and exhibited antiviral activity against Herpes, Measles, Picornavirus and Rubella viruses (Coelho et al., 2014; 2015, 2018). This propolis exhibited low toxicity even at high doses and showed an anti-tumor effect attributed to the inhibition of NO production on Ehrlich tumor developed in mice (Araújo et al., 2011, 2010), beside this, reduced the pathology associated with murine asthma, attributed to an inhibition of inflammatory cells migration to the alveolar space and the systemic progression of the allergic inflammation (de Farias et al., 2014).

Antibiotic-resistant microorganisms have been an ever-growing concern over the past years. Antibiotic resistance occurs when bacteria change in response to the use of medicines. Microbial resistance is a growing threat to the effective treatment of an increasing range of infections, caused by bacteria, parasites, viruses and fungi (WHO, 2014). Since the early 2000s, research for new antibacterial drug, become a long and slow process, characterized by longer development times and lower success rates for investigational drugs (Dheman et al., 2020, Ndagi et al., 2020). Generally, there are not observed differences in antibacterial, antifungal and antiviral activity of propolis from distinct geographic origins, including Brazilian samples from A. mellifera and stingless bees (Sanches et al., 2017). One of the main pharmacological activity of propolis is its ability to inhibit the growth of microorganisms, because its bactericidal and fungicidal functions are essential to preserving life in the hive. Spanish ethanolic extract of propolis containing high amounts of polyphenols exhibited antimicrobial properties (Fernández-Calderón et al., 2020). The presence of pyrrolizidine alkaloids was reported previously in propolis from S. postica (Coelho et al., 2015, 2018). In the present study, the extraction method used allowed the detection of many pyrrolizidine alkaloids, together flavonoids.

Alkaloids are distributed in many classes including pyrrolizidines, pyrrolidines, quinolizidines, indoles, tropanes, piperidines, purines, imidazoles, and isoquinolines. Pyrrolizidine alkaloids occur in several species of plants relevant for human and animal nutrition, being found in the angiosperm family of Boraginaceae (all genera), Asteraceae (tribes Senecioneae and Eupatorieae) and Fabaceae (genus Crotalaria) (Schramm et al., 2019; Moreira et al., 2018; Tasca et al., 2018). These compounds are produced by plants as defense chemicals against herbivores (Schramm et al., 2019; Mädge et al., 2020). Pyrrolizidine alkaloids (PA) and their N-oxides detected in honey and pollen bee products are composed of a 1-hydroxymethylated necine core, which is esterified with a variety of necic acids (Tasca et al., 2018; Kopp et al., 2020; Sixto et al., 2019; Celano et al., 2019, Hungerford et al., 2019, Mädge et al., 2020). Despite the hepatotoxic, genotoxic, cytotoxic, tumorigenic, and neurotoxic activities of 1,2-unsaturated PAs, they can be used for the treatment of diseases and infections, due to their glycosidase inhibitory activity, which led to antidiabetic effect, besides anticancer, fungicidal, and antiviral effects (Schramm et al., 2019; Moreira et al., 2018; Tasca et al., 2018; Mädge et al., 2020).

Scaptotrigona aff. postica collected in Barra do Corda possess potential for chemical and biological exploration, due to its antiviral activity against herpes and rubella viruses, among other diseases (Coelho et al., 2015, 2018; Araújo et al., 2011, 2010). In this study was carried out the characterization of compounds found in the fractions of aqueous extract of propolis (40AEP) and methanolic extract of propolis (40MEP) from S. postica collected in Barra do Corda. Its antimicrobial action was evaluated against strains of fungi, Gram-positive and Gram-negative bacteria, besides the evaluation of the cell viability by the MTT technique.

## Material and methods

Propolis from Scaptotrigona aff. postica was collected in 2017, into the internal parts of a beehive located in the municipality of Barra do Corda (05°30’20”S, 45°14’36”W), state of Maranhão, Brazil, at the confluence of the Rio Corda and Rio Mearim and was kindly provided by Beekeeper Wilson Amorin Melo.

Preparation of the aqueous extract of propolis (40AEP) and methanolic extract of propolis (40MEP)

Generally, the bioactive constituents of propolis are extracted using organic solvents, to eliminate inert materials, while preserving the polyphenolic fractions. Extraction with ethanol is particularly suitable for obtaining deparaffinated extracts rich in polyphenolic components. On the other hand, extraction with pure water is suitable for obtaining extracts containing water-soluble phenolic acids (Kubiliene et al. 2015). Extraction started by macerating of the propolis sample (100 grams) in liquid nitrogen until the obtention of a powder. After this, ultrapure water (Direct-Q 5UV) (100 mL for every 10 grams) and methanol (70%) (100 mL for every 10 grams) was added in different PYREX Erlenmeyer Flask, and was kept under magnetic rotation (IKA® Big Squido - 1,000 rpm) for 24 hours at room temperature, protected from light. After this, the supernatant was centrifuged (Eppendorf® centrifuge 5804R- at 4 ° C for 30 minutes at 15,000 rpm) to separate the precipitate (wax) and subsequently filtered (J. Prolab - qualitative filter paper Ø 15,0 cm). The filtrate was lyophilized (SUPERMODULYO Freezer DRYER Thermo Electron Corporation) to obtain the dry extract. The aqueous extract of propolis (AEP) and methanolic extract of Propolis (MEP) after lyophilization, were dissolved in acidified water (trifluoroacetic acid - 0.05% TFA). For the MEP, 10% DMSO was added to the acidified water. The AEP and MEP were kept at −20 ° C until the experiments were carried out.

### Chromatography separation

The AEP and MEP crude extracts were applied to a Sep-Pak C18 cartridge (Water Associates), previously equilibrated with trifluoroacetic acid - TFA (0.05%). The extracts were eluted in three concentrations of solutions of acetonitrile (ACN) 5%, 40% and 80% acidified with 0.05% of trifluoroacetic acid (TFA). The chromatography separation of chemical constituents was performed using the LC-8A Preparative Liquid Chromatography (Shimadzu), and a preparative reverse phase column Shim-pack ODS PREP-C8 (H) (5μm, 20mm × 25cm) for 70 minutes with a constant flow of 8 mL / min using gradients of ACN / TFA solution (0.05%) (A) and ACN / TFA (0.01%) (B). The running gradient used is 0-10 - 2%, 10-65 - 60%, 65-70 - 2%. Absorbance was monitored at 225nm, at room temperature. The fractions were collected manually. Three fractions of each extract were obtained, which were denominated 5 AEP, 40 AEP, 80 AEP, 5 MEP, 40 MEP and 80 MEP. Only the fractions 40 AEP and 40 MEP demonstrated antimicrobial action against Gram-positive and Gram-negative strains and were selected. The analyses of 40 AEP and 40 MEP fractions were performed using liquid chromatography coupled to mass spectrometry (LC-MS), using a binary UFLC system (20A Prominence, Shimadzu Co., Japan) coupled to the Electrospray - Ion Trap - Time of Flight mass spectrometer (ESI-IT-TOF) LCMS-8030 system (Shimadzu, Kyoto, Japan) consisting of an LC pump (LC-20AD), an autosampler (SIL-20ACHT UFTC), a thermostated column compartment (CTO-20AC), UV–VIS detector (SPD-20A), and a triple quadrupole mass spectrometer (LCMS-8030). The samples were resuspended in water / 0.1% acetic acid and analyzed with a C18 column Discovery C18 (5 μm, 50 mm × 2.1 mm), using (A) acetic acid / water (1: 999) and (B) acetic acid / ACN / Water (1: 900: 99) as solvent, with a constant flow of 0.2 mL / min, where the gradient varied from 5 to 70% of solvent B, during 35 minutes at 37 ° C. The fractions 40 AEP and 40 MEP were monitored at 214 nm by a Shimadzu SPD-M20A PDA detector. The UV spectra of most 1,2-unsaturated PAs show an absorption maximum at about 214 nm (EFSA 2011). The mass spectrometry analyzes were performed, according to the following parameters: the interface voltage used was 4.5 KV and the detector voltage 1.76 KV, with a temperature of 200° C; fragmentation was caused by argon collision gas, with 50% energy; and the spectra were obtained in the range of 50 to 2000 m / z. Argon was used as the nebulizer and desolvation gas. The neutralizer gas flow, drying gas flow, capillary voltage, and nebulizer pressure were set at 3.0 L/min, 15.0 L/min, 6000 V, and 250 °C, respectively. Constituents were detected in ion Trap - Time of Flight mass spectrometer in positive-ion mode. LubSolutions software (Shimadzu, Tokyo, Japan) was used for system control and data analysis. The data obtained were compared to MS data reported by El-Shazly and Wink (2014), Kopp et al., (2020), Mädge et al., (2020), Tian et al., (2017), Cisilotto et al, (2018), Coelho et al., (2015, 2018), Hungerford et al., (2019), Chong and Chua (2020) and Kumar and Chandra (2015).

### Determination of Minimum Inhibitory Concentration (MIC) Microorganisms

Escherichia coli (SBS363), Micrococcus luteus (A270) donated by the Pasteur Institute (Paris, France); Candida albicans (MDM8) donated by the Department of Microbiology at the University of São Paulo (São Paulo, Brazil); Escherichia coli (D31-streptomycin resistant), Staphylococcus aureus (ATCC 29213), Salmonella entrica serovar typhimurium (ATCC 14028) Bacillus megaterium (ATCC 10778) and Micrococcus luteus (BR1 - streptomycin resistant) donated by the American Collection of Cell Culture types (ATCC). Fungi from human clinical isolates Candida krusei (IOC 4559), Candida glabrata (IOC 4565), Candida parapsilosis (IOC 4564), Candida guilliermondii (IOC 4557), Candida tropicallis (IOC 4560) and Candida albicans (IOC 4558) were donated by Oswaldo Cruz Institute (Rio de Janeiro, Brazil). To determine the MIC, the technique of inhibiting microbial growth in liquid medium was used with modifications (Riciluca et al., 2012). The test was carried out in 96-well microplates with a final volume of 100 μL. The samples were evaluated at concentrations of 31.3 to 200 μg / mL, in the culture medium containing 105 CFU / mL for bacteria and / or 104 CFU / mL for fungi. The growth media used were PB (Peptones 10 g / L; NaCl 5 g / L; pH 7.4) for bacteria and PDB (Potato Dextrose Broth; 1.2 g Potato dextrose; 100 mL of H2O; pH 5.0) for fungi. In this experiment, 10 mg / mL positive control and 20 μL ultrapure water (Direct-Q 5UV) were used as a negative control. A solvent control, for all microorganisms, was performed with the maximum concentration of DMSO used to solubilize the samples from the MEP (0.5%). The experiments were carried out in triplicate. Culture growth controls were carried out, as well as appropriate sterility controls of the materials, adding 100 μL of ultrapure water (Direct-Q 5UV) and 100 μL of the culture media. Microbial growth was evaluated by turbidity of the medium by absorbance measurement (λ: 595nm) in a Victor^3^ microplate reader (1420 Multilabel Counter / Victor^3^ - Perkin Elmer), after 24 hours of incubation at 37 ° C.

### Cell viability assay

VERO cells isolated from kidney epithelial cells extracted from an African green monkey Cercopithecus aethiops, were cultured at 37°C in T25 flasks containing 5 mL of Leibovitz medium (L-15) supplemented with 10% fetal bovine serum. After reaching 70% confluence, the cells were treated with trypsin, dissolved in culture medium and counted in a Neubauer chamber, using the trypan blue cell exclusion method (0.2%), in which only viable cells remain discolored. To calculate viability, the following formula was used: % viable cells = number of live cells in the 4 quadrants × 100 total cells counted in the 4 quadrants.

### Cell viability by the MTT technique

The development of the rapid colorimetric assay which relies on the ability of mitochondrial dehydrogenase enzymes to convert 3-(4,5-dimethylthiazol-2-yl)-2,5-diphenyltetrazolium bromide (MTT) to a purple formazan precipitate, has simplified large scale screening of cells. Cell viability was quantified through the activity of succinate dehydrogenase, a mitochondrial enzyme active in cells with intact respiratory chain metabolism, by the MTT method, as described by Mosmann (1983) with modifications.

VERO cells isolated from kidney epithelial cells extracted from an African green monkey Cercopithecus aethiops, were seeded at a concentration of 104 cells / well in 96-well plates in L-15 medium supplemented with 10% fetal bovine serum and maintained at 37 ° C for 24 hours. Upon reaching 70 to 80% of confluence, the wells were washed with sterile PBS to remove non-adherent cells and 100 μL of the samples (at the maximum concentration used in the antimicrobial tests, that is, 200 μg / mL) were added. Positive controls (cells + L-15, 10% SBF) and negative controls (cells + 1% DMSO) were performed. The plates were kept at 37°C, for 24 hours. After 24 h of incubation the culture supernatant was removed and 20 μL of MTT (5 mg / ml) solution in PBS was added to each well. The cells were kept for 4 hours at 37 ° C protected from light. After the period, 100 μl of DMSO was added for 10 minutes for the solubilization of the precipitates. The reading was performed on the ELISA reader (FlexStation® 3, Molecular Devices, USA) at a wavelength of 540 nm. The experiment was carried out in triplicate. The calculations for cell viability values followed the following procedure: 1. Calculation of the average optical density of the samples, cell control and control with 0.5% dimethyl sulfoxide (DMSO). 2. Subtraction of the control mean with DMSO 0.5% of the control means and samples; 3. Calculation of average cell viability using the equation: VC = ODD / ODC × 100; where VC = cell viability; ODD = average of the optical density of the samples; ODC = average of the optical density of the controls. From the data obtained, the graph of cell viability was plotted as a function of the concentration of the samples.

## Results

The extraction method can affect the antimicrobial properties of extracts, extraction yields, as well as the contents of phenolic and flavonoid compounds (Pobiega et al., 2019). The use of UHPLC equipment provided high resolution for the separation of alkaloids and flavonoids and improved the sensitivity of Q-TOF-MS detector. The chromatographic profile of fractions 40 AEP and 40 MEP revealed the presence of 26 compounds, which were tentatively characterized by the interpretation of their mass spectra (MS) compared with MS data reported in the literature, as can be seen in Table 1, including retention times, [M+H]^+^ ions corresponding to protonated molecules and relevant MS/MS ions. Figure 1 illustrates the chromatogram profile of fraction 40 MEP in ESI positive mode. The pyrrolizidine alkaloids (**1-3**, **5**, **7-9**, **13**, **14**, **16**, **18 - 26**), quinolone alkaloids (**4** and **6**), mangiferin (**10** and **17**), flavones glycosides (**11** and **15**) and the tetrahydroxyflavanone aromadendrin-7-O-methyl ether (**12**) were detected in fraction 40 MEP (Figure 1). On the other hand, compounds **1**, **2**, **4**, **6**, **8**, **12** and **21-23** were not detected in fraction 40 AEP.

**Table 1.**
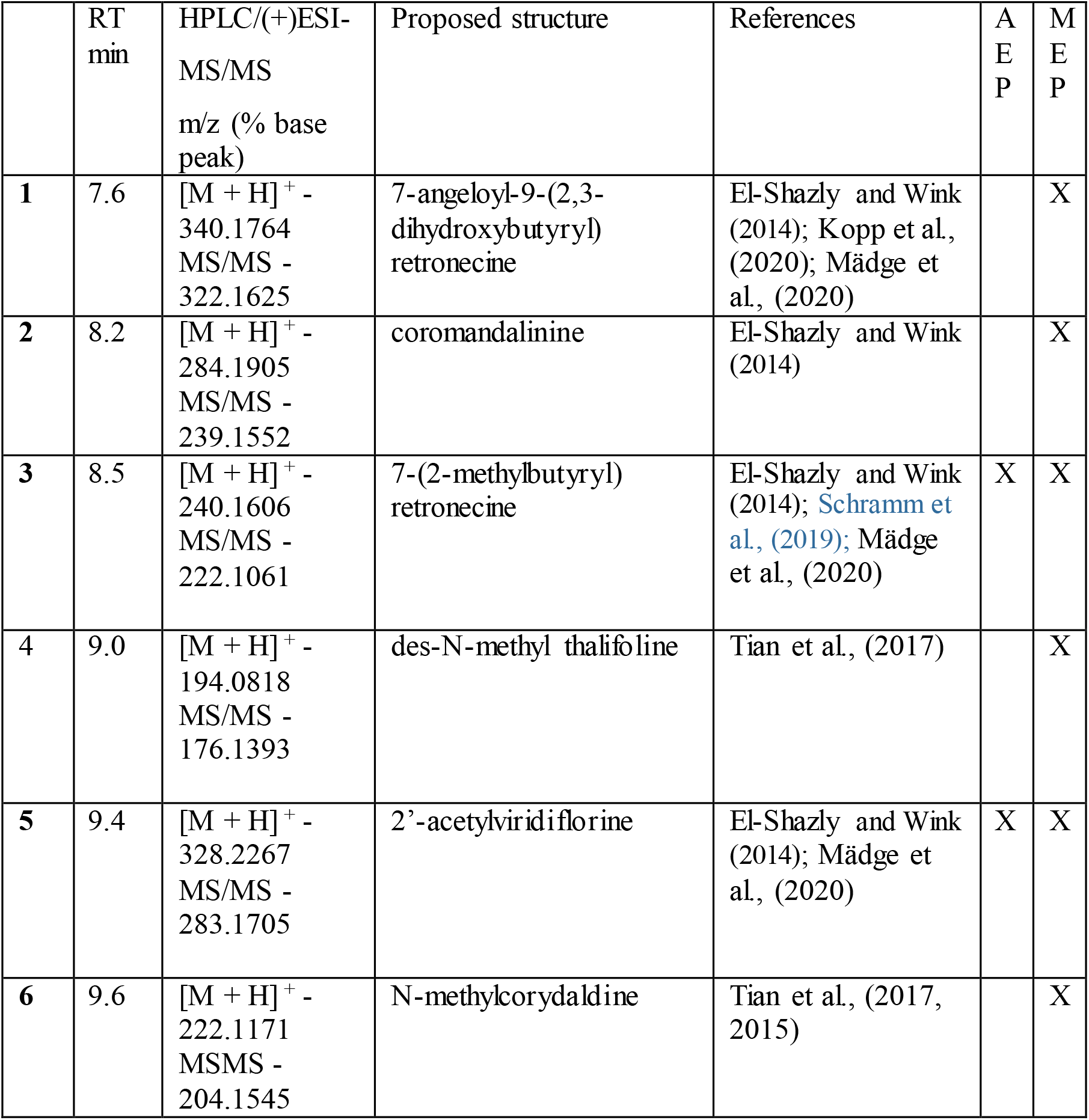
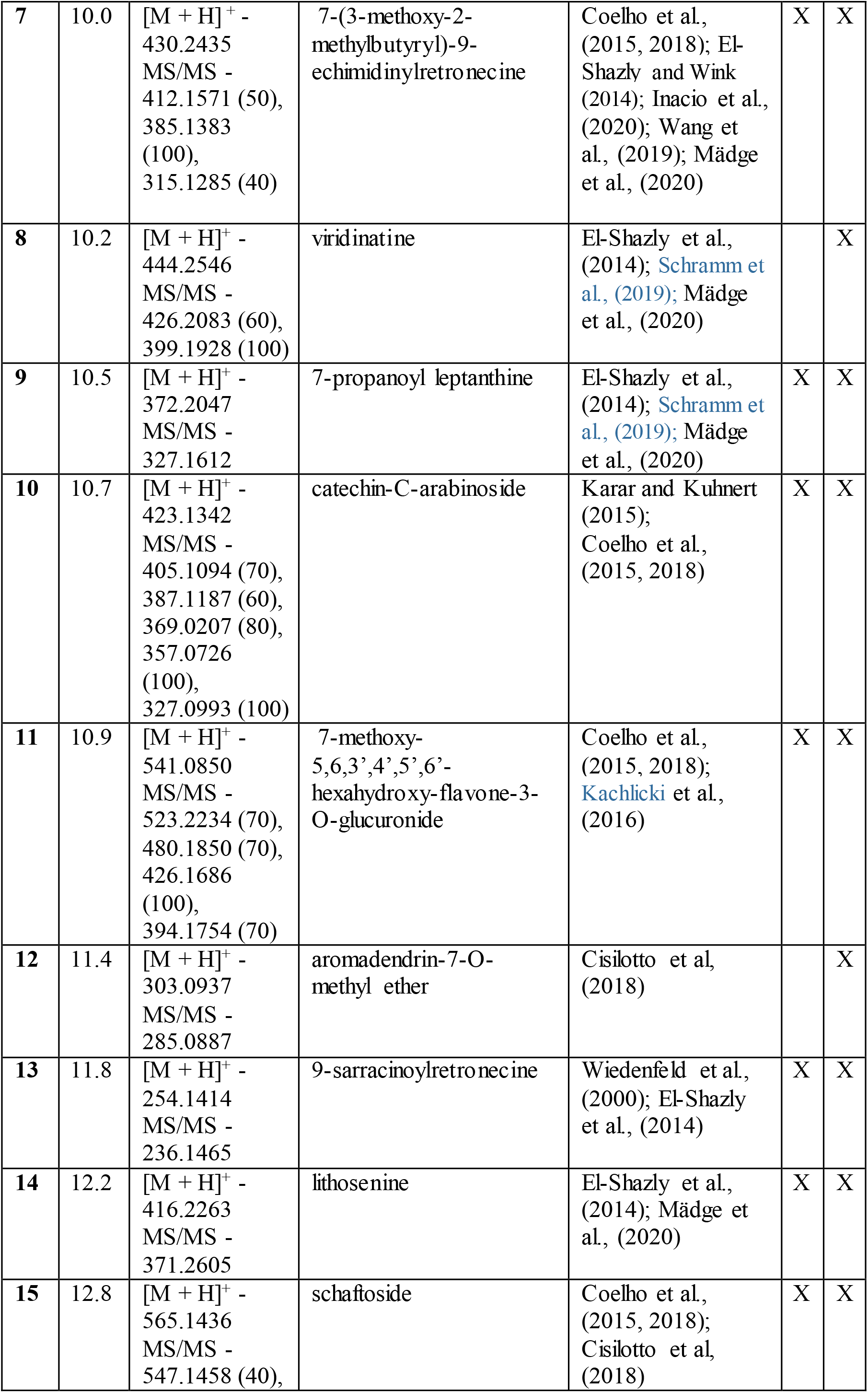
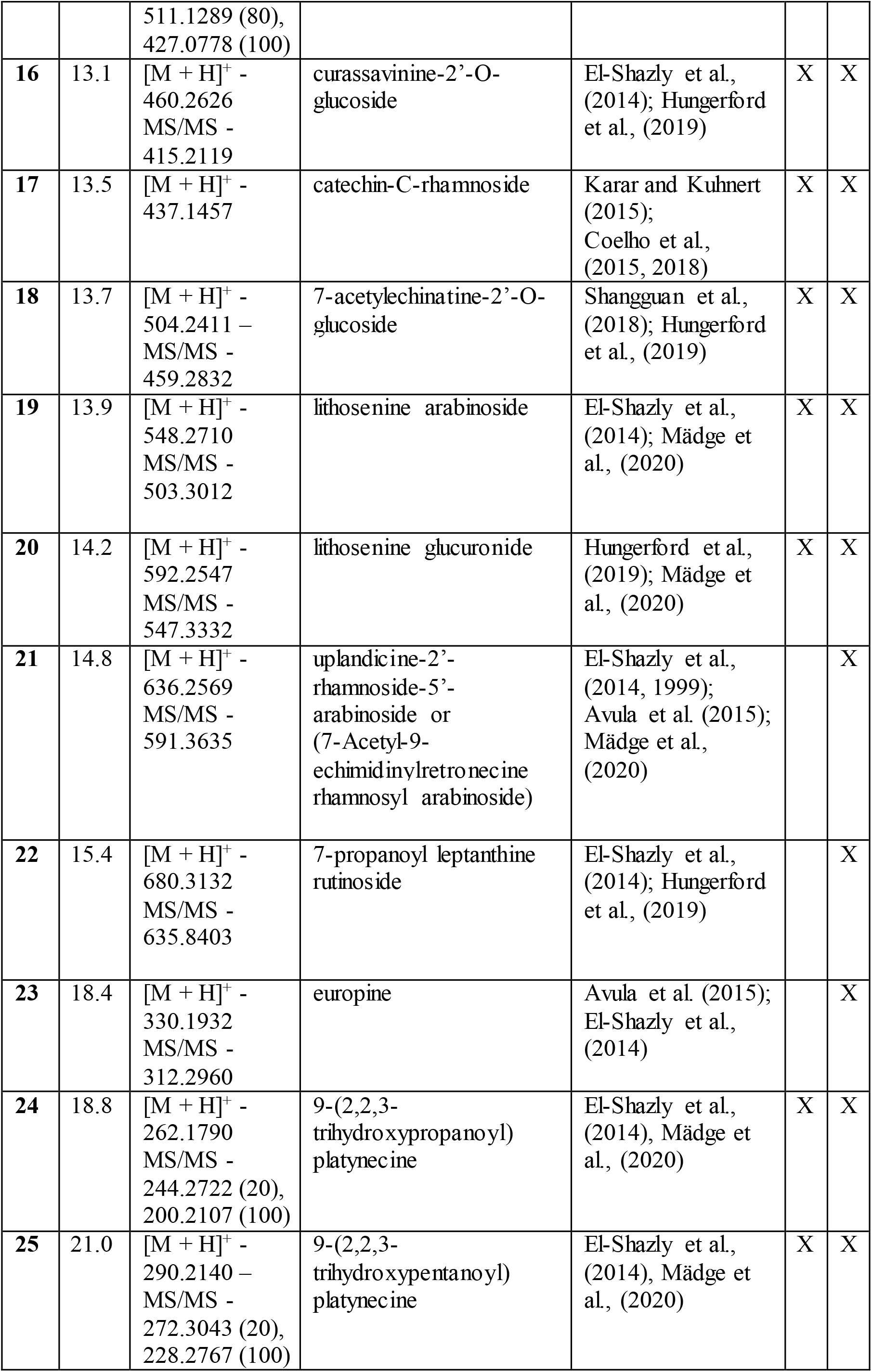
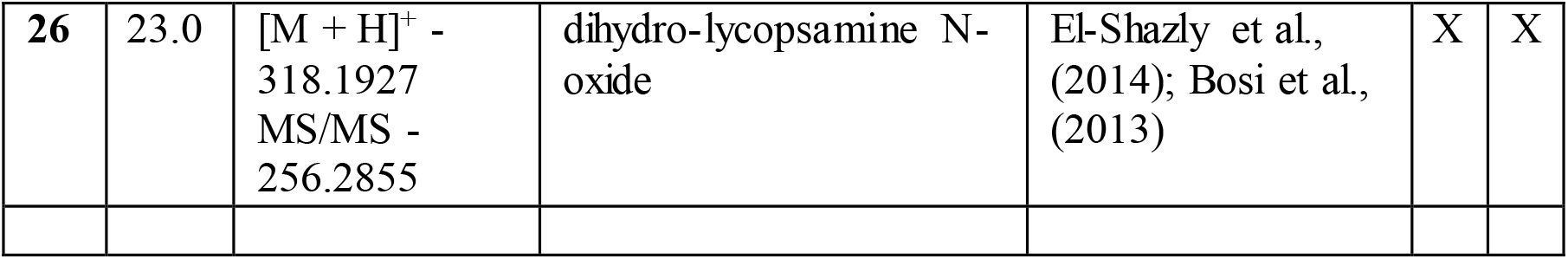
Constituents tentatively characterized, which were detected in aqueous extract of propolis (AEP) and methanolic extract of propolis (MEP) from S*captotrigona* aff. *Postica* RT: retention time; HPLC/(+)ESI-MS/MS mass spectra data in positive ionization mode, proposed structure and literature references.

**Figure 1.**
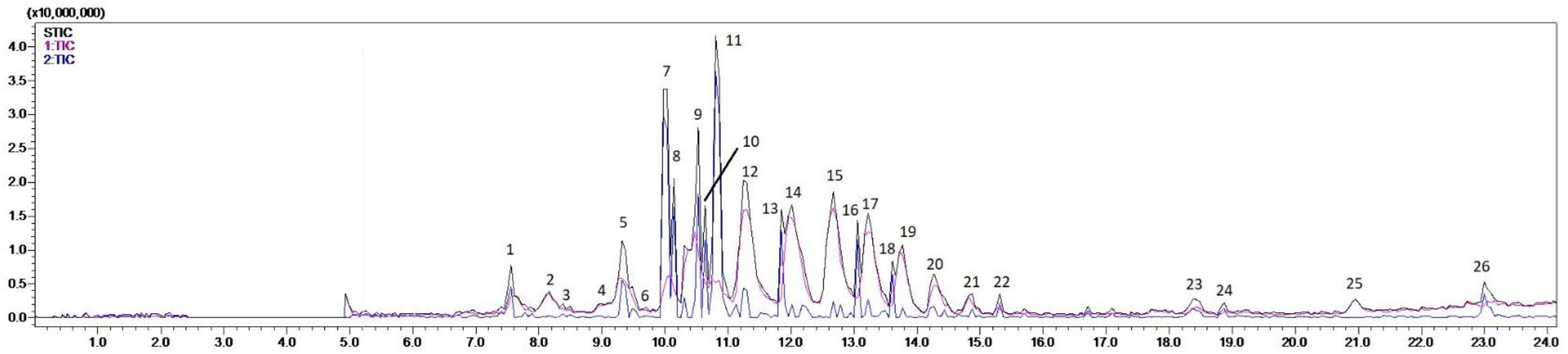
Chromatogram profile of fraction 40 MEP in ESI positive mode by liquid chromatography coupled to mass spectrometry (LC-MS). Source: Image obtained through the LabSolutions software

The chemical structure of pyrrolizidine alkaloids (PA) comprises the necine bases: platynecine, heliotridine, retronecine, and otonecine esterified with one or two of the following necic acids: 2-methyl butyric acid, angelic acid, echimidinic acid, viridifloric acid, trachelanthic acid, sarracinic acid, lasiocarpic acid and curassavic acid, at C-7 and/or C-9 of necine base (Avula et al., 2018; Hungerford et al, 2019; El-Shazly and Wink, 2014; Mädge et al., 2020). Generally, diastereomeric PAs cannot be distinguished based on their MS/MS spectra, because protonated molecule [M+H]+ ions and fragment ions are identical, even with high resolution mass spectrometry (Hungerford et al., 2019). The pyrrolizidine alkaloid, 7-(3-methoxy-2-methylbutyryl)-9-echimidinylretronecine (**7**) exhibited the protonated molecule [M+H]+ with *m/z* 430 and deprotonated molecule [M − H]− at *m/z* 428 detected in both positive and negative ESI mode, respectively. The MS/MS experiments in deprotonated molecule at *m/z* 428, produced abundant fragment ion at **m/z** 397 (echimidine -7-angelyl-9-echimidinylretronecine), attributed to the loss of methoxy (OCH3) group. The MS/MS experiments in protonated molecule at *m/z* 430, produced fragment ions at *m*/*z* 412 attributed to the loss of water (18 Da), base peak at *m/z* 385 produced by the loss of 45 Da, attributed to the loss of (CH3CHOH) group from the echimidinic acid esterified at C9 position of retronecine and the fragment ion [M + H − 115]+ at *m*/*z* 315 was attributed to the loss of methoxy dihydro angelic acid moiety, esterified on the C7 position of the retronecine (Coelho et al., 2015, 2018, Hungerford et al, 2019; ElShazly and Wink, 2014; Mädge et al., 2020). The loss of 45 Da were also observed for the pyrrolizidine alkaloids tentatively assigned as uplandicine-2’-rhamnoside-5’arabinoside (**21**), lithosenine (**14**), lithosenine arabinoside (**19**) and lithosenine glucuronide (**20**), which also possess echimidinic acid esterified at C9 position of retronecine, besides coromandalinine (**2**), 2’-acetylviridiflorine (**5**), viridinatine (**8**) and 7-acetylechinatine-2’-O-glucoside (**18**) which possess viridifloric acid esterified at C9 position of retronecine. For 7-propanoyl leptanthine (**9**) and 7-propanoyl leptanthine rutinoside (**22**); the loss of 45 Da were observed through the loss of (CH_3_CHOH) group from trachelanthic acid esterified at the C9 position of retronecine, while for curassavinine-2’-O-glucoside (**16**) through the loss of (CH3CHOH) group from curassavic acid esterified at the C9 position of retronecine. On the other hand, for 7-angeloyl-9-(2,3-dihydroxybutyryl) retronecine (**1**), 7-(2-methylbutyryl) retronecine (**3**), 9-sarracinoylretronecine (**13**), that possess sarracinic acid (Wiedenfeld et al., 2000), and europine (**23**), that possess lasiocarpic acid esterified at C9 position of retronecine, the base peak was formed by the loss of water (18 Da). The dihydro-lycopsamine N-oxide (**26**) exhibited pseudo-molecular ion [M + H]+ at *m/z* 318 and base peak at *m/z* 256, formed by the loss of 62 Da, corresponding to the loss of water plus carboxy group [M + H − CO2 − H2O]+ (Bosi et al., 2013). For the quinolone alkaloids, compounds **4** and **6**, with pseudo-molecular ion peak [M + H]+ at *m/z* 194 and *m/z* 222, respectively, the base peak was formed by the loss of water (18 Da)(Tian et al., 2017). The platynecine derivatives, 9-(2,2,3-trihydroxypropanoyl) platynecine (**24**) and 9-(2,2,3-trihydroxypentanoyl) platynecine (**25**) exhibited pseudo-molecular ion [M + H]+ at *m/z* 262 and *m/z* 290, respectively, and the base peak were formed by the loss of 62 Da.

Schaftoside (**15**) and 7-methoxy-5,6,3’,4’,5’,6’-hexahydroxy-flavone-3-O-glucuronide (**11**) were reported previously in propolis from S. postica (Coelho et al. 2015, 2018). Aromadendrin-7-O-methyl ether (**12**), exhibited pseudo-molecular ion peak [M + H]+ at *m/z* 303 and base peak at *m/z* 285. This tetrahydroxyflavanone was also detected in propolis from Scaptotrigona bipunctata (Cisilotto et al, 2018), while aromadendrin was detected in geopropolis of Melipona interrupta and M. seminigra (Silva et al., 2013). Mangiferin (**10**) and 3-O-methyl mangiferin (**17**) exhibited protonated precursor ion [M + H]+ at *m/z* 423 and at *m/z* 437, respectively. The MS/MS spectrum of mangiferin (**10**) exhibited fragment ions at *m/z* 405 (100) and *m/z* 387 (80) corresponding to subsequent loss of two water units, while the MS/MS spectrum of 3-O-methyl mangiferin (**17**) exhibited fragment ions at *m/z* 419 (100), and at *m/z* 401 (80) (Kumar and Chandra (2015).

### Determination of Minimum Inhibitory Concentration (MIC)

Minimum inhibitory concentration (MIC) is the lowest concentration of an antimicrobial compound that will inhibit the visible growth of a microorganism after incubation. The 40 MEP showed antibacterial effect against Gram-positive and Gram-negative bacterial pathogens with MIC ranging from 62.5 μg / mL to 200 μg / mL (Table 2). The highest antibacterial activity was observed against Bacillus megaterium ATCC10778. On the other hand, 40 AEP showed antibacterial effect against Gram-positive bacterial pathogens with MIC of 200 μg / mL and showed inactivity against Gram-negative bacteria, as can be seen in Table 2. MEP and AEP were inactive against fungi Candida genus, which cause the most common fungal infections worldwide.

**Table 2.** Determination of the Minimum Inhibitory Concentration (MIC) of the crude aqueous (AEP) and methanolic (MEP) extracts

| Microrganisms | 40 AEP | 40 MEP |
| --- | --- | --- |
| **Gram-positive bacteria** |  |  |
| *Staphylococcus aureus* ATCC 29213 | ND | **200** |
| *Bacillus megaterium* ATCC10778 | **200** | **62.5** |
| *Micrococcus luteus* A270 | **200** | **125** |
| **Gram-negative bacteria** |  |  |
| *Escherichia coli* SBS 363 | ND | **200** |
| *Escherichia coli* D31- streptomycin resistant | ND | **200** |
| **Fungi** |  |  |
| *Candida krusei* IOC 455916 | ND | ND |
| *Candida parapsilosis* IOC 456416 | ND | ND |
| *Candida guilliermondii* IOC 4557 | ND | ND |
| *Candida albicans* IOC 45588 | ND | ND |
| *Candida tropicalis* IOC 45608 | ND | ND |
| *Candida glabrata* IOC 45658 | ND | ND |

MIC was expressed in µg / mL. The tested concentrations were from 31.1 to 200 µg / mL. (ND) Did not show antimicrobial activity.

### Assessment of cell viability

Cytotoxicity and biocompatibility tests simulate biological reactions to bioactive molecules when they are placed in contact with cells and body tissues. The use of cell lines can predict whether a molecule might be toxic to the body, since it is possible to measure how much the cell metabolism was affected during the test. Although tests assessing cytotoxicity in vitro may not have correlation with in vivo tests, if a propolis extract induces a proven cytotoxic reaction in cell culture tests, it is much likely that it will develop toxicity when used in foods, treatment of diseases and cosmetics (Utispan et al., 2017; Saavedra et al., 2016). The cell viability test was carried out using 3- (4,5-dimethylthiazol-2-yl)-2,5- diphenyl tetrazolium bromide (MTT) reduction assay. The cells were put in contact with 40MEP and 40AEP at a concentration of 200 μg / mL, and their viability was assessed after 24 h by reaction with the MTT reagent. As the 40MEP was resuspended with 0.5% DMSO, a control was performed with DMSO in the same concentrations to verify its interference in cell growth. Cell viability was expressed in terms of the relative absorbance of treated and untreated cells (control). The result of the effects of the concentrations of 40AEP and 40MEP, on the activity of succinate dehydrogenase were expressed in percentage of cell viability as can be seen in Figure 2. This Figure shows the relative viability of fibroblast culture of Cercopithecus aethiops and was not observed variations with respect to control cells. Thus, no significant changes were observed at 200 μg / mL of 40AEP and 40MEP. There was no statistical difference between treated and untreated cells.

**Figure 2.**
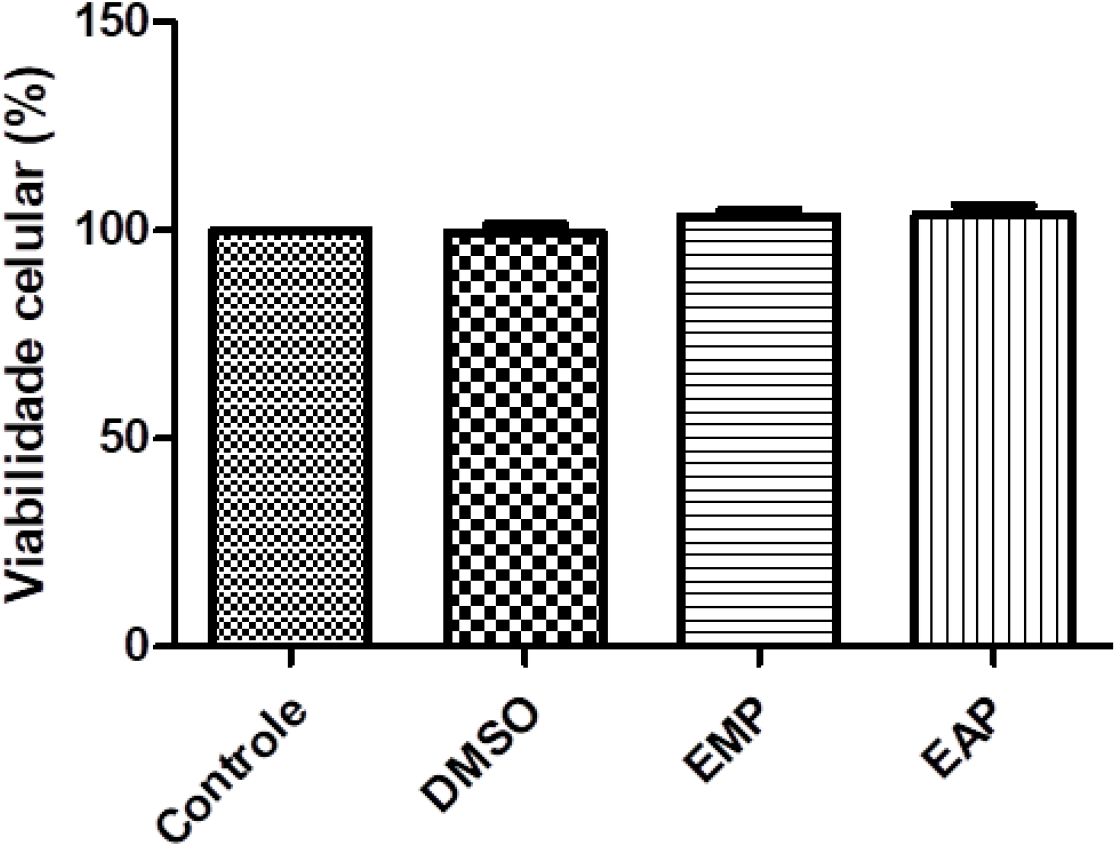
Treatment with AEP and MEP samples, from propolis from S. aff. postica on cell viability of fibroblast culture after 24h of treatment. Cell viability was assessed using the MTT colorimetric assay. Source: ANOVA GraphPad Prism 5.1.

## Discussion

### Chemical composition

Although studies about propolis or geopropolis of stingless bees indicated that they are chemically similar with propolis from Apis mellifera, some chemical characteristics observed in propolis of meliponine differ from the chemical profile of propolis from honeybees. Propolis or geopropolis from stingless bee are richer in flavonoid glycosides (Kachlicki et al., 2016) than Apis mellifera propolis. Propolis of A. mellifera contains larger amounts of flavonoid aglycones and the scarce contents of glycosides. This occur due to a direct consequence of the presence of the enzyme β-glucosidase, which through enzymatic process enables the rapid hydrolysis of the flavonoid glycosides from resin source during propolis collection and processing (Zhang et al., 2012). The chemical profile of propolis from S. postica is interesting and different. The presence of a pyrrolizidine alkaloids was reported previously (Coelho et al., 2015, 2018). However, in this sample were found a high content and variety of pyrrolizidine alkaloids, as can be seen in Table 1. The presence of alkaloids was reported also for a sample of geopropolis from Scaptotrigona bipunctata bees collected in the state of Santa Catarina (South Brazil) (Cisilotto et al, 2018).

The beekeeping products, honey, pollen, royal jelly, propolis, and beeswax, are vulnerable to PA contamination, attributed to nectar and plant pollen rich in pyrrolizidine alkaloids (PAs) collected by bees and transferred to bee products (Brugnerotto et al., 2021, Sixto et al., 2019, Celano et al., 2019, Wang et al. 2019, De Jesus Inacio et al., 2020). Lycopsamine-type PAs/PANOs were predominant in bee pollen, while heliotrine-type PAs/PANOs were less frequent and contributed with lower contents (De Jesus Inacio et al., 2020). Echimidine, lycopsamine, and intermedine were the most abundant alkaloids detected in honey, along with lesser amounts of each of their N-oxides (Kowalczyk et al., 2018, Hungerford et al., 2019, Wang et al., 2019). Echimidine-N-oxide was the main alkaloid detected in honey and nectar samples, while echivulgarine-N-oxide was the main PA found in plant pollen, indicating that nectar contributes more significantly to PA contamination in honey than pollen collected in plant (Lucchetti et al., 2016, Kast et al., 2019, Picron et al., 2019, Gottschalk et al., 2020). 3’-O-Glucosyllycopsamine and 3’-O-glucosylintermedine and the corresponding N-oxides were identified in honey samples containing high contents of lycopsamine-type PAs/PANOs (Hungerford et al., 2019). Toxicity of the PAs involves unsaturation of the 1,2 position and esterification of at least one of the hydroxyl groups with an acid and increases with the degree of branching and formation of long cyclic diesters. However, PAs are pro-toxins as they require metabolic activation to exert toxic effects (Sixto et al., 2019, Celano et al., 2019, Wang et al. 2019, De Jesus Inacio et al., 2020, Hungerford et al., 2019).

### Antimicrobial activity

Propolis was approved as an official drug in the London pharmacopoeia in the 17th century, due to its antibacterial properties (Przybyłek and Karpiński, 2019; Okińczyca et al., 2020, Hochheim et al., 2019, 2020). The antimicrobial activity of propolis and geopropolis was demonstrated in clinical, in vivo, and in vitro studies (Cornara et al., 2017, Pasupuleti et al., 2017, Siheri et al., 2016). Other bee products also exhibited antimicrobial activity. Honey from stingless bees, Scaptotrigona bipunctata Lepeletier, and S. postica Latreille, exhibited in vitro antimicrobial activity against Gram-positive and Gram-negative bacteria, including multidrug-resistant strains (Nishio et al., 2016).

Propolis and its constituents exhibited activity against targets in the pathophysiological context of the disease caused by SARS-CoV-2 (Berretta et al., 2020). Propolis exhibited anti-bacterial activity against Gram-positive and Gram-negative, as well as aerobic and anaerobic bacteria (Przybyłek and Karpiński, 2019; Okińczyca et al., 2020) and exhibited antifungal activity against pathogenic yeasts C. albicans, C. parapsilosis, C. tropicalis, and C. glabrata (Cornara et al., 2017, Przybyłek and Karpiński, 2019; Siheri et al., 2016). Although, there are many studies about the antimicrobial properties of propolis, the comparison of the studies is difficult, due to the varying composition of propolis extracts and the different methods used for the evaluation their antimicrobial activity (Pasupuleti et al., 2017; Okińczyca et al., 2020; Torres et al., 2018). Poplar propolis exhibited antimicrobial and antiviral activities, besides anti-inflammatory activities (Governa et al., 2019). Extracts of red, green, and brown propolis from different regions of Brazil, demonstrated antimicrobial activity against Staphylococcus aureus (Dantas Silva et al., 2017). The ethanol extract of red propolis from Alagoas exhibited antimicrobial activity against Staphylococcus aureus and Candida krusei (Silva et al., 2019), and antifungal activity against resistant fungal isolates of C. parapsilosis and C. glabrata (Pippi et al., 2015). Albanian propolis inhibited both microbial growth and biofilm formation of Pseudomonas aeruginosa, an opportunistic pathogen responsible for a wide range of clinical conditions (Meto et al., 2020). Propolis extracts from various geographic origins (Germany, Ireland, and Czech Republic) synergistically enhanced the efficacy of antibiotics, vancomycin and oxacillin, against drug-resistant microorganisms (Al-Ani et al., 2018).

Brazilian and Venezuelan propolis from stingless bees, Melipona quadrifasciata, Melipona compresssipes, Tetragonisca angustula, and Nannotrigona sp. exhibited high antibacterial activity (Dos Santos et al., 2017b). The hydroalcoholic extract of propolis from Melipona quadrifasciata showed high antibacterial activity against S. aureus and P. aeruginosa and significant antimollicute s activity against Mycoplasma pneumoniae (Hochhein et al., 2020, Dos Santos et al., 2017b). Geopropolis from Heterotrigona itama showed antibacterial activity against Gram-positive (Bacillus subtilis and Staphylococcus aureus) and Gram-negative bacteria (Escherichia coli and Pseudomonas aeruginosa) (Abdullah et al., 2019). Geopropolis from Melipona mondury exhibited bacteriostatic and bactericidal activity against Pseudomonas aeruginosa, Staphylococcus aureus, and methicillin resistant Staphylococcus aureus (Dos Santos et al., 2017c). The extract of geopropolis from Melipona quadrifasciata quadrifasciata was effective against S. aureus and methicillin-resistant S. aureus (Torres et al., 2018). Propolis from stingless bees Tetragonisca fiebrigi exhibited antimicrobial activity against gram-positive and gram-negative bacteria and fungi (Campos et al., 2015).

The results of this study are corroborated by many studies, in which were observed that propolis acted more efficaciously against Gram-positive than Gram-negative bacteria. The structure of Gram-positive bacteria cell wall, with predominant share of peptidoglycan, allows hydrophobic molecules to penetrate the cells and act on wall as well as cell membrane and within the cytoplasm. The cell wall of Gram-negative bacteria is more complex with less peptidoglycan and with outer membrane composed of double layer of phospholipids linked with inner membrane by lipopolysaccharides (Ristivojević et al., 2016), beside this, can produce a wide range of hydrolytic enzymes. Thus, Gram-negative may be more resistant to active constituents of propolis (Ristivojević et al., 2016). The antiviral activity of propolis can be attributed to the cell lysis and disruption of the cytoplasmic membrane of virus leading to leakage of cellular components and eventually cell death (Górniak et al., 2019, Tagousop et al., 2018, Coelho et al., 2015, 2018). Studies demonstrated that propolis has different antibacterial mechanisms, including inhibition of cell division, collapsing microbial cytoplasm cell membranes and cell walls, inhibition of bacterial motility, different bacterial enzymes activity, what can be cause the weakens the stability of cytoplasmatic membrane (Okińczyca et al., 2020) and acting in bacteriolysis and protein synthesis inhibition (AlAni et al., 2018, Aboody and Mickymaray, 2020, Górniak et al., 2019, Tagousop et al., 2018). Although some studies demonstrated antifungical activity of propolis against Candida genus, this propolis sample did not exhibited antifungical activity this fungus.

A cell viability assay is performed based on the ratio of live and dead cells. This assay is based on an analysis of cell viability in cell culture for evaluating in vitro extracts effects in cell-mediated cytotoxicity assays for monitoring cell proliferation (Saavedra et al., 2016; Utispan et al., 2017). In the present work was possible to measure how much the cell metabolism was affected by the 40MEP and 40AEP extracts and the results of cell viability demonstrated that these extracts were not toxic to fibroblast culture of Cercopithecus aethiops. Reactive oxygen (ROS) and nitrogen (RNS) species serve as molecular signals activating stress responses that are beneficial to the organism, however, can cause oxidative damage and tissue dysfunction (Saavedra et al., 2016; Utispan et al., 2017). The huge interest in the evaluation of cell viability after exposure to extracts, is important due to the constituents presents in extracts, which can act in the prevention of ROS and RNS formation and exhibit protective effects against cancer and inflammatory diseases (De Francisco et al., 2018).

Cellular toxicity of Chilean propolis extract was evaluated using MTT and apoptosis/necrosis detection assays and the results demonstrated the role of polyphenols in the control of extracellular matrix degradation in atherosclerotic plaques (Saavedra et al., 2016). Propolis from Trigona sirindhornae, a new species of stingless bee, differentially inhibit the proliferation of a metastatic neck squamous cell carcinoma cell line (Utispan et al., 2017). The oral usage of propolis extract from S. postica exhibited low toxicity, even at high doses (Araújo et al., 2011). In this study, using an in vitro model of Vero cells isolated from kidney epithelial cells extracted from an African green monkey Cercopithecus aethiops, the treatment with 40MEP and 40AEP from propolis in concentrations 200 μg / mL did not interfere in cell viability. Pyrrolizidine alkaloids are considered toxic, however, in this study, the results of cell viability demonstrated that these compounds did not cause cytotoxicity in epithelial cells extracted from an African green monkey Cercopithecus aethiops.

Infections caused by multidrug-resistant microorganisms are associated with increased mortality compared to those caused by susceptible bacteria and they carry an important economic burden. Multidrug resistant bacteria can compromise the clinical utility of major chemotherapeutic antimicrobial agents. Antimicrobial compounds, as flavonoids found in propolis, exhibit a broad spectrum against different types of bacteria and can enhance the efficacy of conventional antibiotics (Almuhayawi, 2020). Flavonoids are used for many centuries in the treatment of the range of human diseases due to nontoxic effect (Górniak et al., 2019, Tagousop et al., 2018, Adamczak et al., 2019). Schaftoside was found in high content in 40MEP and 40AEP. Flavonoid-C-glycosides exhibited antiviral, antibacterial and antifungal activities (Xiao et al., 2017). The presence and position of the sugar group in the flavone glycosides generally, had no effect on the MIC values (Adamczak et al., 2020). Flavonoids demonstrated more potent activity against Gram-negative bacteria: E. coli and P. aeruginosa than Grampositive ones: E. faecalis and S. aureus (Górniak et al., 2019, Tagousop et al., 2018, Adamczak et al., 2019).

Most alkaloids are found to be bactericidal rather than being bacteriostatic. The alkaloids mechanism of action as antibacterial agents is different for each type of alkaloid (Thawabteh et al., 2019). Alkaloids can form hydrogen bonds with enzymes, receptors, and proteins, due to a proton accepting nitrogen atom. Thus, alkaloids can be able of inhibiting the activity of bacteria, fungi, protozoan and other microorganisms (Thawabteh et al., 2019, Othman et al., 2019). Pyrrolizidine alkaloids (1-3, 5, 7-9, 13, 14, 16, 18 – 26) and quinolone alkaloids (4 and 6), were detected in fraction 40 MEP. Pyrrolizidine alkaloids (PA) exhibited antioxidant and antimicrobial activity in vivo and in vitro, affecting cell division, respiratory inhibition enzyme, inhibition in bacteria, bacterial membrane disruption and affecting virulence genes (Othman et al., 2019, Belyagoubi-Benhammou et al., 2019, Schramm et al., 2019). The growth of bacterial species, mostly human pathogens, such as E. coli, Streptococcus pneumoniae, B. subtilis, Bacillus anthracis and S. aureus, was inhibited in the presence of different pure PAs and plant extracts containing PA (Joosten and van Veen, 2011). Pyrrolizidine alkaloids interacted with human immunodeficiency virus (HIV) activity (Schramm et al., 2019, Moreira et al., 2018) and acted as promising prototypes for new drugs especially for topical use (da Silva Negreiros Neto et al., 2016). Pyrrolizidine alkaloids can inhibit protozoan parasites, such as Plasmodium, Trypanosoma, Leishmania, Trichomonas and intestinal worms exhibiting antiparasitic properties, through typical DNA damage, that occurs when DNA alkylating compounds, such as, pyrrolizidine alkaloids form covalent bonds with DNA bases (Wink, 2012).

## Conclusion

Emerging bacterial resistance is a major concern for the medical field. Each propolis sample has its own chromatographic profile and biological activity, which is related to its botanical origin. Schaftoside and pyrrolizidine alkaloids were detected in MEP and AEP, which exhibited antibacterial activity, mainly against Gram-positive bacteria. It is difficult to compare the results of different studies, due to the varying composition of propolis extracts and the different methods used for the extractions of constituents and evaluation their antimicrobial activity. The results of this study showed that the biological properties depend on factors, such as the extraction method and region from which propolis was collected. In this study the extraction method allowed the detection of new pyrrolizidine alkaloids, which are considered toxic, however demonstrate pharmacological properties. Pyrrolizidine alkaloids can be exploited by relying in medicinal chemistry strategies that can maintain bioactivity while reducing toxicity.

## Acknowledgment

Research support Foundation of State of São Paulo (FAPESP/CeTICS), Grants n° 2013/07467-1 and Brazilian National Council for Scientific and Technological Development (CNPq) Grant n° 422744/2012-7

